# A single-cell transcriptomic landscape of innate and adaptive intratumoral immunity in triple negative breast cancer during chemo- and immunotherapies

**DOI:** 10.1101/2021.12.01.470716

**Authors:** Laura Carpen, Paolo Falvo, Stefania Orecchioni, Giulia Mitola, Roman Hillje, Saveria Mazzara, Patrizia Mancuso, Stefano Pileri, Alessandro Raveane, Francesco Bertolini

**Affiliations:** Laboratory of Hematology-Oncology, European Institute of Oncology IRCCS, 20141 Milan, Italy; Human Technopole, 20157, Milan, Italy; The Hyve, Utrecht, Netherlands; Hematopathology Unit, European Institute of Oncology IRCCS, 20141 Milan, Italy

## Abstract

Breast cancer (BC) constitutes a major health problem worldwide, making it the most common malignancy in women. Current treatment options for BC depend primarily on histological type, molecular markers, clinical aggressiveness and stage of disease. Immunotherapy, such as anti-PD-1, have shown combinatorial clinical activity with chemotherapy in triple negative breast cancer (TNBC) delineating some therapeutic combinations as more effective than others. However, a clear overview of the main immune cell populations involved in these treatments has never been provided.

Here, an assessment of the immune landscape in the tumour microenvironment (TME) of two TNBC mouse models (4T1 and EMT6 cell lines) has been performed using single-cell RNA sequencing (scRNA-seq) technology. Specifically, immune cells were evaluated in untreated conditions and after being treated with chemotherapy or immunotherapy used as single agents or in combination. A decrease of regulatory T cells, compared to the untreated TME, was found in treatments with *in vivo* efficacy as well as γδ T cells, which have a pro-tumoral activity in mice. Focusing on Cd8 T cells, across all the conditions, a general increase of exhausted-like Cd8 T cells was confirmed in pre-clinical treatments with low efficacy; on the other hand, an opposite trend was found for the proliferative Cd8 T cells. Regarding macrophages, M2-like cells were found enriched in treatments with low efficacy while opposite behaviour was associated with M1-like macrophages. For both cell lines, similar proportions of B cells were detected with an increase of proliferative B cells in treatments that involved cisplatin in combination with anti-PD-1. The fine-scale characterization of the immune TME in this work can lead to new insights on the diagnosis and treatment of TNBC for a possible application at the clinical level.

## Introduction

Breast cancer (BC) is the most common malignancy in women, and one of the three most common cancers worldwide, along with lung and colorectal cancer making it the fifth leading cancer death worldwide ^1^. Histological and molecular analysis have confirmed that BC is a heterogeneous disease, and among its subtypes, the triple negative breast cancer (TNBC) accounts for 15-20% displaying the worst prognosis in terms of disease-free survival and overall survival ^2^. TNBCs are predominantly undifferentiated, with high proliferative capacity and metastatic potential ^2^. Gene-profiling studies have confirmed the heterogeneity of this disease and have led to a better understanding of its molecular profile with the identification of new possible therapeutic targets ^3,4^.

Until recently, the backbone of therapy against TNBC has been chemotherapy; that differs in mechanisms of action in alkylating, such as cyclophosphamide ^5^ and cisplatin ^6^; antimicrotubule such as taxanes ^7^ and antineoplastic agents such as doxorubicin ^8^. Cancer immunotherapy has changed the landscape of cancer treatment during the past few decades, highlighting the importance of the interaction between the immune system and cancer ^9^. These therapies have different approaches that include for example the use of specific drugs such as monoclonal antibodies which can be small proteins or fusion proteins that act on the surface of cancer or immune cells. Immune checkpoint inhibitors (ICI) are the most clinically used approaches and act by controlling the activation and the intensity of the adaptive immune response. Their activity is crucial to avoid exacerbated immune responses and autoimmunity by the induction of T lymphocyte exhaustion^10^. The two immune checkpoint receptors that have been most studied in the context of clinical cancer immunotherapy are cytotoxic T-lymphocyte-associated antigen 4 (CTLA-4) and programmed cell death protein 1 (PD-1) ^11 12–14^.

PD-1 predominantly regulates effector T cell activity within tissue and tumours by binding the programmed cell death ligand 1 (PD-L1). In turn, this binding inhibits kinases involved in T cell activation ^15^. In physiological conditions, the interaction of PD-1 with its ligands has been shown to play an important role in the maintenance of the balance between autoimmunity and peripheral tolerance ^16^, serving as an immunological regulator and to maintain an immune homeostasis ^17^. In the tumor microenvironment (TME), PD-1 and its ligand PD-L1 perform a vital role in progression and survival of cancer; the overexpression of PD-L1 by tumour cells is used as self-defence by the tumour against the cytotoxic T cells which contribute to cell killing ^18^. PD-L1 expression on many tumours is a component of a suppressive microenvironment that leads to T cell dysfunction and exhaustion ^19^. This state of exhaustion is characterised by the progressive loss of proinflammatory cytokines production, the loss of the cytotoxic activity, the decrease in the proliferative potential and an increase in apoptosis ^20^. As a consequence, blocking the PD-1/PD-L1 inhibitory pathway can re-activate T cells in the TME with the release of inflammatory cytokines and cytotoxic granules to eliminate tumour cells. PD-1 is also highly expressed on regulatory T cells (Tregs), where it may enhance their proliferation in the presence of PD-L1 ^21^. Because many tumours are highly infiltrated with Tregs that suppress effector immune responses, blockade of the PD-1 pathway may enhance antitumor immune responses by diminishing the suppressive activity of intratumoral Treg cells.

Breast cancer has been considered an immune silent cancer type that is less likely to benefit from immunotherapy. However, among BC subtypes, TNBC is believed to be a more immunogenic subtype and therefore potentially responsive to ICIs ^22^. Anti-PD-1 and anti PD-L1 have shown combinatorial clinical activity with chemotherapy in TNBC, but only in a minority of patients and only for a limited period of time. Moreover, it is currently unclear which cell populations are involved in the immune response within the TME during specific conditions as well as their proportion in specific treatments ^23^.

Single-cell RNA sequencing (scRNA seq) gives the possibility to differentiate among cell populations that are not distinguishable by cell surface markers and morphology alone, opening the possibility of identifying previously uncharacterised cellular populations, phenotypes and transitional states. The advancements of this technique have revolutionized our ability to study the immune system and allow us to break through the bottleneck of immunology studies ^24^, providing valuable resources for both basic research and clinical applications ^25^.

In this work, we have investigated at the single cell level, report and discuss in detail the transcriptome of innate and adaptive intratumoral immune cells in two syngeneic, immune competent, orthotopic murine models of local and metastatic TNBC. Mice were treated with ICIs and several different types of chemotherapeutics, alone or in combination. From previous reports, capecitabine (alone or with ICIs) was the less effective drug ^26^. While platinum, doxorubicin and taxanes showed synergy with ICIs and had superimposable activity, intermittent, medium dosage cyclophosphamide (C140) plus vinorelbine and ICIs was the most active combinatorial therapy. Vinorelbine activated antigen presenting cells and C140 generated new T cell clones including stem cell-like TCF1+ CD8+ T cells ^27^. The fine characterization of almost 50,000 immune cells extracted from the TME of these two mouse models helped in creating a catalogue of the immune response to several drugs and aimed to investigate specific cellular subtypes useful for future therapeutic approaches.

## Materials and Method

### Cell lines and treatments

As this work focuses on the computational analysis of scRNAseq data previously published, *in vivo* and *in vitro* experiments were performed as mentioned in ^27^. Briefly the laboratory procedures included the injection of two TNBC cell lines (4T1 and EMT6) in the mammary fat pad as in ^26^. Tumour-bearing mice were treated with either vehicle or with different drugs used as single agents or in combination as described in ^27^ for a total of eight treatments and one untreated control for each cell line (Table 1). Drug usages were based on literature data associated with no or acceptable toxicity ^27,28^.

**Table 1.** Schematic view of conditions (samples) with the corresponding treatment for the 4T1 cell line

| Label | Type of treatment | Chemistry 10X<br>for 4T1 | Chemistry 10X<br>for EMT6 |
| --- | --- | --- | --- |
| ctr | control | v2 | v2 |
| C140<br>(Cyclophosphamide) | alkylating agents | v3 | v3 |
| C140 + V<br>(Vinorelbine) | alkylating agents +<br>anti-microtubule<br>agents | v3 | v3 |
| C140 + αPD-1 | alkylating agents +<br>ICI | v2 | v2 |
| C140 + V + αPD-1 | alkylating agents +<br>anti-microtubule<br>agents + ICI | v3 | v2 |
| D (Doxorubicin) +<br>αPD-1 | antineoplastic agents<br>+<br>ICI | v2 | v3 |
| P (Cisplatin) +<br>αPD-1 | antineoplastic agents<br>+ ICI | v2 | v2 |
| T (Paclitaxel) +<br>αPD-1 | anti-microtubule<br>agents +<br>ICI | v3 | v2 |
| αPD-1 | immuncheckpoint<br>inhibitor (ICI) | v2 | v2 |

### ScRNA library preparation and sequencing

At 28 or 70 days, depending on the efficacy of the treatment and on the cell line (details in ^27^), tumour resection was performed as in ^28^.

Tumours of three mice were dissociated and pooled together to generate the single cell suspension. The cell suspensions were then prepared for cell sorting with FACS Fusion sorter (BD bioscience). Only Cd45 + DAPI - (alive immune cells) were sorted and purity evaluated; at least 5,000 cells per condition underwent scRNA-seq library preparation following the 10X Genomic protocol and using two different chemistries (v2 and v3) (Table 1). The sequencing was performed with NovaSeq^™^ 6000 Illumina^®^ sequencer at a sequencing depth of 50,000 reads pairs/cell.

### Alignment and quality control

FASTQ files were converted to digital gene-cell count matrices using a Singularity-dependent Snakemake pipeline ^29^ employing the Cell Ranger v4 software. As reference, the *Mus musculus* reference genome mm10 (GENCODE vM23/Ensembl98) was used. Market Exchange Format (MEX) for sparse matrices generated from the pipeline were loaded, merged, processed and analyzed using Seurat package v4 ^30^.

To have a comparable number of cells for each experiment, we excluded from the analyses all the conditions with a number of cells lower than 500. Then, in order to include only cells that are of highquality, we exclude cells with 500 or less transcript, 50,000 or more transcripts, having fewer than 250 expressed genes, a complexity score (log10 genes per UMI) lower than 0.80 and more than 15% mitochondrial transcripts as in ^27^. At the gene-level, all genes expressed in less than five cells were filtered out.

To detect doublets, the *scDblFinder()* function of scDblFinder package ^31^ was used. This package works only with SingleCellExperiment (SCE) objects ^32^, therefore, a conversion from Seurat object was performed. Finally, doublets were excluded using the *subset()*function of the Seurat package^30^.

During the QC, one condition (C140+V+αPD-1), out of the nine reported here for the 4T1 cell line, was removed from further analyses because its number of cells did not exceed the minimum filtering threshold (500 cells) used by us to consider a condition suitable to explore the whole immune populations. For the EMT6, all conditions passed this quality control filter. The number of transcripts and number of genes were evaluated for the remaining conditions. The majority of the cells had more than 1,000 UMI indicating high quality cells (Fig. S1). Furthermore, in the 4T1, the C140, C140+V and T+αPD-1 treatments had the highest number of mitochondrial transcripts; while for the EMT6 it was C140+V, C140+V+αPD-1 and P+α-PD-1. This might be related to differences in toxicity of these specific agents in different microenvironments ^33–35^

### Proliferation status, normalization and batch effect removing

Cell cycle score variation was evaluated with a Principal Component Analysis (PCA) on normalized and scaled data using 2,000 genes on Seurat v4 package after assigning a cell cycle score to each cell with the *CellCycleScoring()* function and the S and G2M specific gene reference downloaded from the *Ensemldb* R package ^36^. No large differences were observed among cell cycle phases between the two cell line TMEs, therefore, we did not regress out the cell cycle variation in the following normalization step.

A batch effect is an unwanted source of variation resulting in different cells having specific profiles, not because of their biological features but because of technical differences. Our data presented a strong batch effect due to the two different types of chemistry used during the single cell library preparations (Fig. S2A). Recently, Seurat introduced the scRNA-seq integration workflow, a set of methods to match shared cell populations across different batches ^30^. These methods identify cross-batch pairs of cells that are in a matched biological state (‘anchors’). In detail, we applied the integration workflow that included splitting of the raw transcript count matrix by chemistry, normalization using the *SCTransform()* function, selection of the most variable features (genes) using *SelectIntegrationFeature()* function, preparation of the Seurat object for the interrogations with *PrepSCTIntegration()* function, canonical correlation analysis (CCA) with *FindIntegrationAnchors()* function and final integration across conditions with *IntegrateData()*. These steps corrected the unwanted source of variations (Fig. S2B) and were applied before each clustering analysis.

### Dimensionality reduction, visualization and clustering

The Uniform Manifold Approximation and Projection (UMAP) method for visualization was employed on the first 30 principal components using the *RunUMAP()* function of the Seurat package. In order to identify known (previously identified cell populations) or uncharacterised cell types, the Seurat v4 graph-based clustering approach, which exploits a K-nearest neighbour (KNN) graph, was applied. We determined the k-nearest neighbour graph with *FindNeighbors()* function and then performed the clustering with the *FindClusters()* function from resolution 0.4 to 1 in steps of 0.2. The resolution 0.8 and 0.6 were evaluated as best for the 4T1 and EMT6 TMEs respectively, according to the number of cell populations (clusters) that were possible to detect. The choice of the best granularity parameters was evaluated by visual inspection and with the aid of the *Clustree* package ^37^.

### Cell-type annotation

The *SingleR* package ^38^ in combination with the *ImmGen()* reference transcriptome dataset ^39^, containing 253 fine labels generated from 830 microarray samples of sorted cell populations, was used for automatic cell type assignment. We inspected the confidence of the predicted labels using the delta values: the difference between the score for the assigned label and the median across all labels for each cell. Using the *PruneScores()* function, we marked potentially poor-quality or ambiguous assignments based on the delta value. Moreover, we uniformed the label name of the *ImmGen()* dataset according to the wanted level of resolution by using the cell ontology label present in the celldex package ^38^. For example, two of the several Cd4 T cell labels were T.CD4.24H (CL:0000624) and T.CD4.CTR (CL:0000624), therefore, we searched for the cell ontology label in Ontology Lookup Service (OLS) repository (https://www.ebi.ac.uk/ols/ontologies/cl) and established “T cells Cd4” as a common label. We verified the assignment using two procedures: i) exploring the expression of known cell gene markers; ii) evaluating the top differential expressed genes (DEG) between cell clusters on *PanglaoDB* ^40^.

Differentially expressed genes were retrieved using the *FindAllMarkers()* function in the Seurat package with a MAST test ^41^. Only genes expressed on 25% of cells and with a log fold-change higher than 1.5 were considered. For this analysis, the normalized data (not integrated) was used as suggested by the Seurat developers (https://github.com/satijalab/seurat/issues/2014#issuecomment-629358390).

Beside using Seurat, DEG analysis was also performed using the Single-Cell Experiment R package (SCE) ^32^ that, differently from Seurat, allows a block on batch. This block, necessary in our dataset, would reveal biologically relevant genes to be preserved within the batch (Table S1, 4T1; Table S2, EMT6).

A label name, mirroring the cell composition, was assigned to each set of cells under the same group (cluster). If multiple cell populations were present in a cluster the first name refers to the most abundant type of cell.

### Sub-clustering

Cells in the macro-clusters of interest (Cd4 T cells, Cd8 T cells and NK cells, Macrophages and B cells) were extracted according to their label name using the *subset()* function in Seurat package. The filtered transcript counts were re-normalized as before using the integration workflow or the classic normalization depending on the purpose of the analysis.

Principal Component Analysis and UMAP methods were applied as before, with the only difference being that we evaluated the best number of PCs to use for the clustering workflow with the *maxLikGlobalDimEst()* function of the *intrinsicDimension* package^42^ as used in ^43^.

Clustering was performed as above and the best resolution was evaluated with the *Clustree* package ^37^. Also depending on the results and the expression of known gene markers the granularity was chosen accordingly (0.2 and 0.3 for Cd4 and Cd8 respectively in 4T1 cell line, 0.1 and 0.2 for B in the 4T1 and EMT6 cell line respectively, 0.4 for macrophages in EMT6 cell line). The sub-clusters cell assignment was performed only with manual curation by choosing a known set of genes from relevant studies that focus on the same cell populations ^44–47^ in similar mouse models and evaluating their expression in the sub-clusters.

### Trajectory analysis

Dynamic changes in gene expression were evaluated by performing a trajectory analysis using the Slingshot package ^48^. To give a finer definition of cell states and unknown cell populations the trajectory analyses were performed only on the cluster subsets.

The *Slingshot()* function was used on the Seurat object converted into SCE dataset ^32^, then the embedding of trajectory in new space was performed with the *embedCurves()* function and finally the *slingCurves()* assessed each curve in each sub-clustering.

## Results

### Total immune cellular landscape in the tumor microenvironment of two TNBC mouse cell lines

After the QC, the resulting total number of cells and genes for the 4T1 were 22,403 and 18,124 respectively, while for the EMT6, 26,245 cells and 18,637 genes were obtained (Table S3, Table S4). The treatments having the highest number of cells after the QC corresponded to T+αPD-1, C140 and C140+αPD-1 for the 4T1; while the C140+V, P+αPD-1 and D+αPD-1 were the treatments with the highest values in EMT6 cell line. Therefore, the analyses focused on a total of 48,648 immune cells and 17 conditions in the TMEs of two TNBC mouse cell lines.

The best granularity resolution (see materials and methods) in the two TMEs identified a total of 20 and 22 groups of cells sharing similar gene expression, for the 4T1 and EMT6 respectively (Fig. 1, Fig. S3). Clusters 0, 13, 15, 18 and 19, in the 4T1 cell line, had the majority of cells automatically assigned (see materials and methods) to B cell population; clusters 1, 2, 3, 4, 6, 7, 8 and 11 contained a high percentage of cells belonging to T cells; cells in clusters 9 and 10 were assigned to macrophages; finally, cluster 5 contained a high percentage of neutrophils, while cluster 14 presented a high percentage of NK cells. Low percentages of unassigned cells were observed in clusters 12, 15 and 19, and very few cells in some clusters were assigned to cell populations that were Cd45^−^ cells such as fibroblast and epithelial cells (Fig. S3A). These together with a need of validation were the reasons for the manual exploration (see materials and methods).

**Figure 1.**
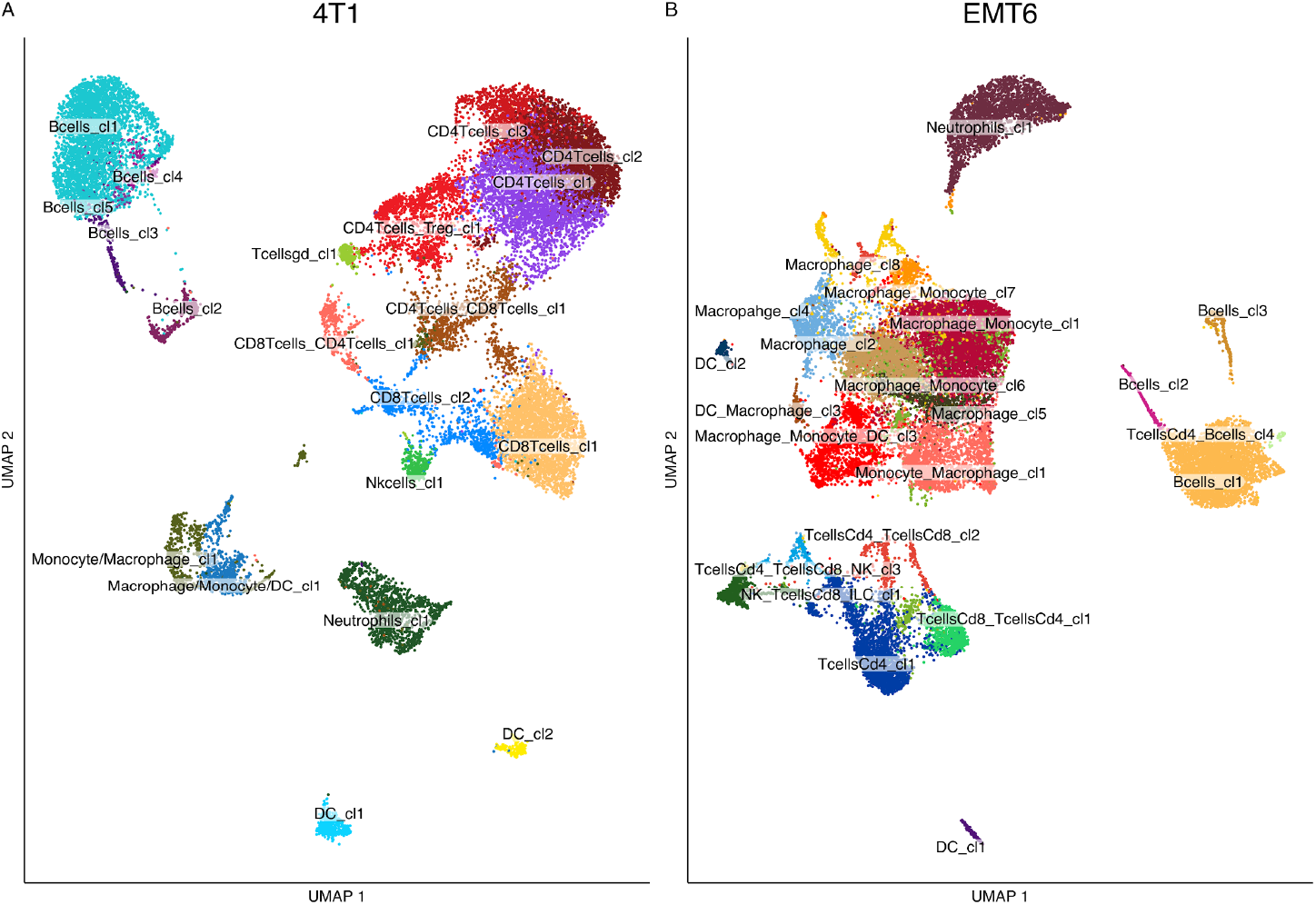
Immune cellular landscape in the two mouse TNBC TMEs.

A) UMAP of 22,403 immune cells in 4T1 TME grouped in 20 clusters. Each dot refers to a cell. The colours refer to the label assigned to the clusters. B) UMAP of 26,245 immune cells in EMT6 TME grouped in 22 clusters. Each dot refers to a cell. The colours refer to the label assigned to the clusters.

EMT6 automatic cell assignment (Fig. S3B) identified clusters 0, 4, 7, 8, 9, 12 and 13 as macrophages; the 3, 10, 14 and 18 clusters were mainly composed of T cell populations; while cells in clusters 2 and 5 were assigned to monocytes. Finally, clusters 1, 15 and 16 were composed mainly by B cells. Compared to 4T1, a higher percentage of unassigned cells was present in EMT6; specifically in clusters 9, 13, 18 and 21. Likely, this was associated with a low expression of gene signature markers revealed by SingleR.

Subsequently, for both the cell lines, *Cd3e*, *Cd4*, *Cd8b1* genes were manually evaluated for T cells, *Ncr1* for NK cells, *Cd19* for B cells, *Csf3r* for neutrophils, *Adgre1* and *Cd68* for macrophages and *Basp1* for DC (Fig. S3C, D). The known gene marker expression in each cluster was in accordance with the highest frequencies of cell populations automatically assigned to that cluster. An additional confirmation was obtained by analysing DEG for each cluster compared to all the others (Table S1, TableS2). These genes were further evaluated on PangloaDB and once again, the results confirmed the automatic assignment.

Final label assignment resulted in the 20 4T1 clusters being classified in 5 B, 5 Cd4 T, 3 Cd8 T, 1 NK, 1 T γδ, 2 DC, 2 macrophage and 1 neutrophil cell clusters (Fig. 1A). On the other hand, the EMT6 cell line presented 22 clusters labelled as 8 macrophage, 1 monocyte, 1 neutrophil, 3 B, 3 DC, 4 Cd4 T, 1 Cd8 T and 1 NK cell clusters (Fig. 1B).

At first glance, a strong difference in the immune cell population composition was found among the two tumoral cell line TMEs. Specifically, the 4T1 TME recorded a prevalence of cells belonging to the lymphoid lineage: 15 clusters contained T, NK and B cells while only 5 were named as macrophages, DCs and neutrophils. On the other hand, in the EMT6 TMEs, most of the cells fall into myeloid clusters while only few cells were assigned to lymphoid lineage. Interestingly, among the lymphoid lineage the number of cells belonging to B cell and neutrophil clusters were comparable between the two tumor types.

Since a fine-scale characterization of the immune landscape wanted to be reached, a filtering and a new sub-clustering for the major immune components of the lymphoid and myeloid lineages, where a informative number of cells could be retrieved, was performed. Due to the differences found in cell population composition, here will be reported the results of Cd4, regulatory, γδ, Cd8 T and NK cells sub-clustering for the 4T1 cell line TME; while of macrophages for the EMT6. Moreover, a comparison of the equally represented B cell clusters between the two types of tumor is also presented and discussed.

### Cd4 T cell-like sub-clustering in 4T1 TMEs reveals pro-tumoral activity of mouse specific T cell population

Cells belonging to clusters that mainly contained Cd4, regulatory and γδ T cells, based on the label assignment, were retrieved and re-clustered (see materials and methods section). The procedure resulted in 7 clusters (Fig. 2A) which can be presented as follows: two progenitor-like Cd4 T cell clusters (0 and 2), characterised by the high expression of *Sell*, *Ccr7*, *Lef1* and *Tcf7* genes (Fig. 2B). A Cd8-like cluster (cluster 3) showing high expression of *Cd8a* gene, these cells are likely a subset of cells deriving from a cluster presenting a mixed cell composition and retrieved because included Cd4 cell population; a T reg-like cell cluster (cluster 1) that presented a high level of *Foxp3* and *Ikzf2* genes; a γδ T cell cluster (cluster 4) that expressed the *Trdc* gene; and finally, two exhausted-like Cd4 T cell clusters (5 and 6) that had a high expression of *Nr4a1* and *Tox* gene markers. Among them, cluster 5 exhibited a more active profile due the high expression of *Cd40Ig* gene but at the same time a higher expression of *Il7r* gene that encodes for a receptor whose ligand was related to tumour progression in γδ T cells ^49^.

**Figure 2.**
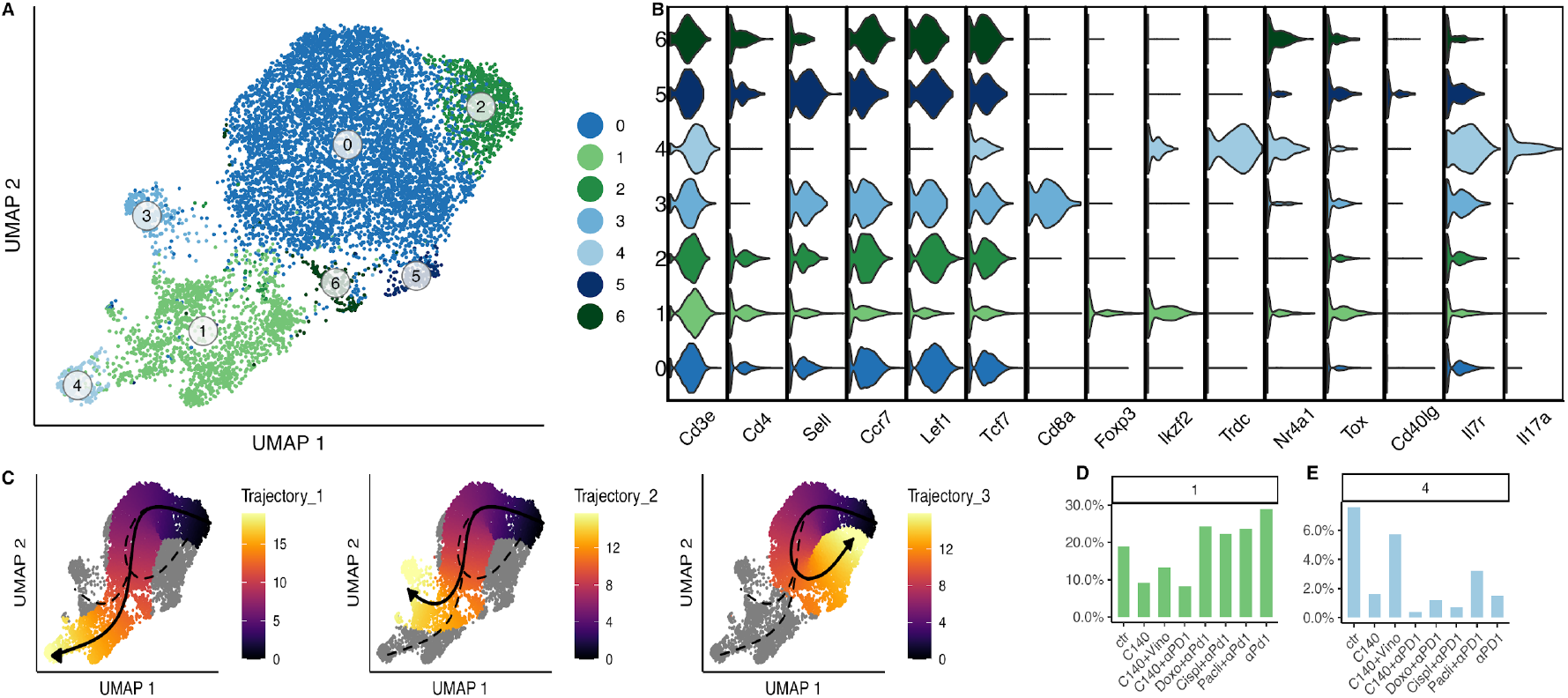
Cd4-like T cell sub-clustering analysis in 4T1.

A) UMAP of Cd4-like T cell sub-clustering in 4T1 cell line TMEs. Each dot refers to a cell. The colours refer to the sub-clusters. B) Gene signature: the violin plot shows the gene on the x-axis, while the clusters on the y-axis. The colours indicate the corresponding cluster. C) Pseudotime trajectory: each dot represents a cell colour-coded for pseudotime. D-E) Proportions of cells in Cd4 T cell sub-cluster among different conditions. Bar graph shows on the x-axis the conditions, while the percentage of cells per cluster is plotted on the y-axis.

UMAP analyses can be used to project the information related to multi-branched trajectories in order to facilitate pseudo-time analysis that measures the relative progression of each of the cells along a biological process of interest without explicit time-series data ^50–52^.

Computationally imputed pseudotime trajectory confirmed and extended the understanding of this sub-cluster composition (Fig. 2C). As previously reported ^53^, three distinct trajectories or cluster differentiations always starting from the same root (cluster 2) were found. The first connects the root (cluster 2) to the γδ T cell cluster passing by the second progenitor-like cluster (cluster 0). The end of the trajectory in the second trajectory is the Cd8-like cluster (cluster 3). While in the third, cluster 2 generates exhausted-like cluster 5 passing through the other exhausted state (cluster 6) (Fig. 2C).

The percentage of cells for each cluster varied significantly among the different conditions (Fig. 2D, Fig. S4). Among the most relevant results, a decrease, compared to the control TME, in the percentage of cells belonging to T reg cluster (cluster 1) was observed in treatments with high in vivo efficacy such as C140, C140+V, C140+αPD-1 ^27^ confirming their immunosuppressive activity. γδ T cell cluster (cluster 4) followed a similar pattern, being high in the control and lower than the control in all the treatments (Fig. 2E). This might be associated with a recently discovered mouse pro-tumoral activity of these specific cell populations ^49,54^. Also confirmed by the high expression of *Il17a*, a marker proved to promote the expansion of pro-tumoral γδ T cells55. The two exhausted-like T cell clusters, related in the pseudotime trajectory analysis, had opposite trends with the more exhausted (cluster 6) being more expanded in treatments with high efficacy than cluster 5 (Fig. S4).

### The Cd8 T cell-like composition in the TME of 4T1 TNBC

We carried out a sub-clustering of Cd8-like T cells like as we did for the Cd4; this led to the identification of 7 clusters (Fig. 3A) that can be associated, also in this case, to different cell types on the basis of a series of gene makers (Fig. 3B). Two progenitor-like Cd8 clusters (0 and 1) were identified as confirmed by the expression of *Sell*, *Lef1*, *Tcf7* and *Ccr7* genes. Three clusters (clusters 3, 4 and 6) had terminally differentiated profiles. Specifically, cells belonging to cluster 3 showed high expression of genes associated with proliferation of phase 2 Cell Cycle such as *Ccnb2*, *Cdk1*, *Mki67* and *Top2a*. Cluster 4 was the most active cluster since it presented high levels of *Gzmb*, *Gzmk*, *Ifng* and *Ly6c1* genes, markers characteristic of effector cells. Furthermore, cluster 6 was defined as exhausted-like Cd8 cluster since it had high levels of *Pdcd1*, *Lag3*, *Ctla4*, *Havcr2* gene expression. Finally, cluster 5 was associated with NKs as the high level of *Ncr1* gene expression suggested (Fig. 3B).

**Figure 3.**
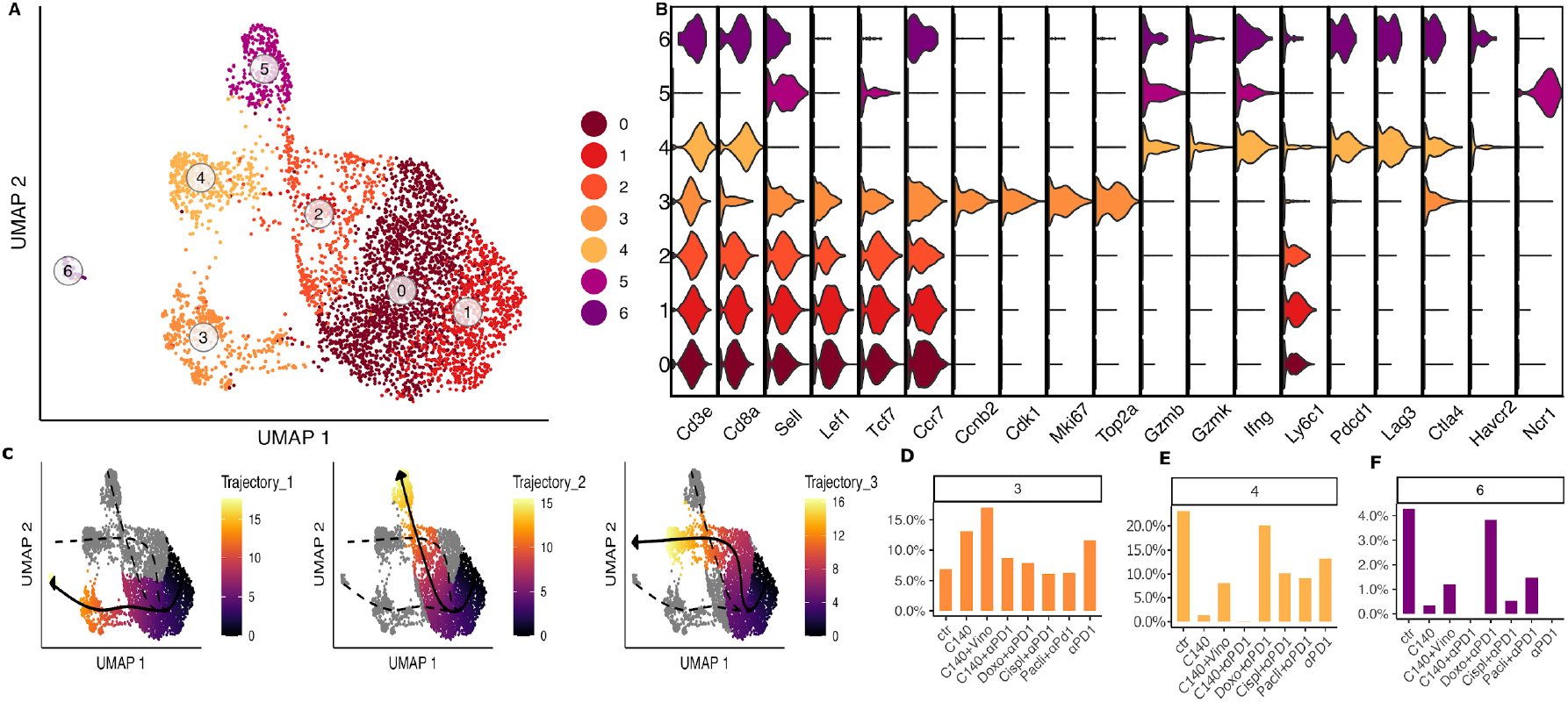
Cd8-like T cell sub-clustering analysis in 4T1.

A) UMAP of Cd8 T cell sub-clustering in 4T1 cell line. Each dot refers to a cell. The colours refer to the sub-clusters. B) Gene signature: the violin plot shows the genes on the x-axis, while the clusters on the y-axis. The colours indicate the corresponding cluster. C) Pseudotime trajectory: each dot represents a cell colour-coded for pseudotime. D-F) Proportions of cells in each 4T1 Cd8-like T cell sub-cluster among different conditions. Bar graph shows on the x-axis the conditions, while the percentage of cells per cluster is plotted on the y-axis.

Looking at the sub-cluster cell proportions along the conditions (Fig. S5), one of the most relevant results was that the highly proliferative Cd8 T cells belonging to cluster 3 increased in TMEs treated with C140 and almost doubled their percentage, compared to the control, in C140 alone and C140+V treated TMEs, underlining a possible anti-tumoral effect. An opposite trend was recorded for the exhausted-like Cd8 T cluster (cluster 6): their cells decreased or were not found in treatments that involved cyclophosphamide while they were present in high percentage in the untreated TME. Similarly, cluster 4 was found enriched in the untreated control while its percentage of cells dropped down in C140, C140+V and C140+αPD-1 treatments (Fig. 3D-F).

Additionally, phenotypic heterogeneity along the Cd8 T cell-like sub-clusters, as for that of the Cd4 cells, was visualized using trajectory analyses. As for the Cd4 sub-clustering, the trajectory of Cd8 sub-clusters (Fig. 3C) revealed three distinct lineages that share a common cluster as root. This cluster was the progenitor-like cell cluster 1. The first trajectory linked the root with the most-exhausted Cd8 cluster (cluster 6). The second related cluster 1 with the cluster referred to NKs; while the third showed a connection with the most-active Cd8 cluster (cluster 4) passing through cluster 2 as for the second trajectory. This analysis confirms both the assignment done previously and the gene expression signature for each cluster.

### M1- and M2 -like tumor associated macrophages populations in the EMT6 TMEs

Abundance in macrophages found in EMT6 TMEs allowed a sub-clustering of these cells. This led to the identification of 9 clusters of cells sharing similar transcriptional profiles (Fig. 4A); of which clusters 2, 5 and 6 were associated to a M2-macrophage subtype because of the expression of *Cx3cr1* gene and the negative expression of *Ly6c1* gene ^56^ (Fig. 4B). Specifically, cluster 6 presented moderate expression of the *Arg2* gene and less extent of *Ptgs2* gene that results in immune suppression ^57^. Instead, cells belonging to cluster 5, highly expressed gene markers associated with proliferation such as *Ccnb2*, *Mki67* and *Top2a* genes defining this cluster as M2-like proliferative. While cluster 2 highly expressed *C1qa/C1qc* genes that were found upregulated in a previously reported M2-like tumor associated macrophage cluster ^47^.

**Figure 4.**
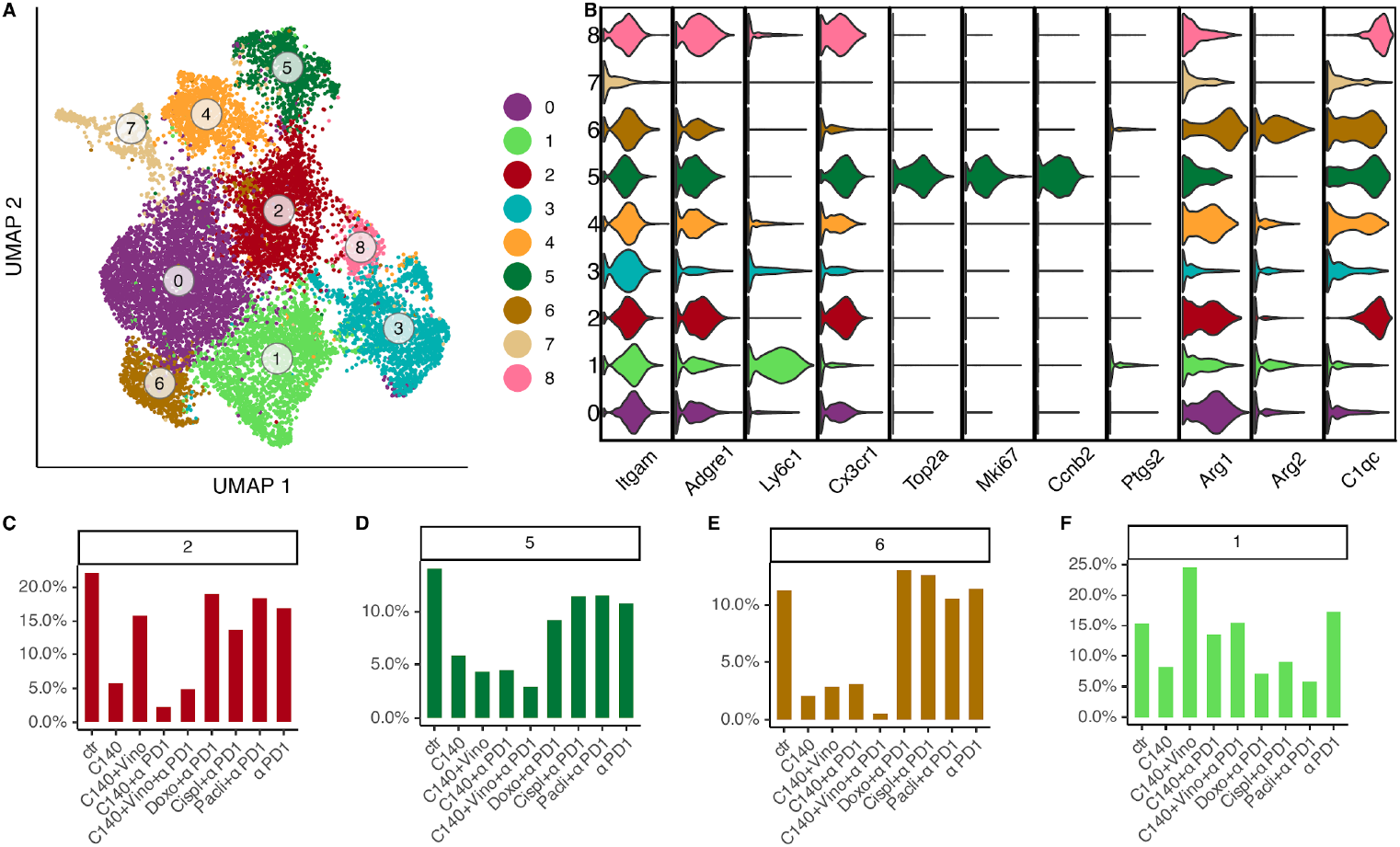
Macrophage-like cell sub-clustering analysis in EMT6.

A) UMAP of macrophages sub-clustering in EMT6 cell line. Each dot refers to a cell and the colours refer to the sub-clusters. B) Gene signature: the violin plot shows the genes on the x-axis, while the clusters on the y-axis. The colours indicate the corresponding cluster. C-F) Proportions of cells in each EMT6 macrophages sub-cluster among different conditions. Bar graph shows on the x-axis the conditions, while the percentage of cells per cluster is plotted on the y-axis.

On the other hand, cluster 1 resulted in being associated with a M1-like macrophagic population due to the expression of *Ly6c1/Ly6c2* gene ^56,58^. The remaining clusters, due to the continuous expression of most gene markers along multiple cell populations, were of uncertain classification as also observed in ^59^ and ^60^.

Observing the proportion of cells for each cluster among the different conditions, the most relevant results were shown when observing the M2-like clusters 2, 5 and 6 (Fig. 4C-E, Fig. S6). They exhibited a similar trend; indeed, their number of cells had high frequencies in the untreated condition and decreased in all the treatments involving cyclophosphamide. On the other hand, M1-like cluster 1 exhibited a remarkable increase only with cyclophosphamide combined with vinorelbine (C140+V) treatment (Fig. 4F).

### Common B cell sub-cluster proportions on both the cell line TMEs

Thanks to the comparable number of cells classified as B cells in the two TNBC cell line TMEs, we explored similarities and differences of the sub-clustering of these cell populations (Fig. 5). The TMEs of the 4T1 cell line displayed five clusters, while eight were the clusters observed in the TMEs of EMT6. Strikingly, the EMT6 model had a more structured sub-clustering revealing more sub-populations. Among these, EMT6 cluster 4 resembled a interferon-induced naïve B cell (expression of *Ifit3*) and clusters 1 and 2, as part of cells belonging to cluster 1 in the 4T1, were classified as an intermediated state between proliferative and naïve-like B cells as clear by the expression of *Pim1* ^61^ (Fig. 5 C,D). On the other hand, clusters 2 and 3, in 4T1 and EMT6 respectively, had a similar transcriptional profile attributable to a proliferative B cell population, as confirmed by the expression of *Mki67* and *Mcm5* genes ^44,62^. Similarly, cluster 4 and cluster 7, in 4T1 and EMT6 respectively, highly expressed *Cd38* gene but not *Mki67* (Fig. 5C,D). These genes have been associated with germinal B cells. Notably, one cluster on the 4T1 (cluster 3) displayed a gene signature typical of plasma B cells (high expression of *Cd27*, *Cd38*, *Xbp1*) ^44,45^ that was not observed in the EMT6 counterpart despite the higher number of subpopulations retrieved. As reported from the scRNA analysis of human nasopharyngeal carcinoma TMEs^30^, a large number of less differentiated B cell subpopulations identified as naïve-like cells were found on both the cell line TMEs. This was the case of cluster 0 for both 4T1 and EMT6, characterized by the expression of *Ighm*, *Ighd* but not *Cd2 7* ^30,45^.

**Figure 5.**
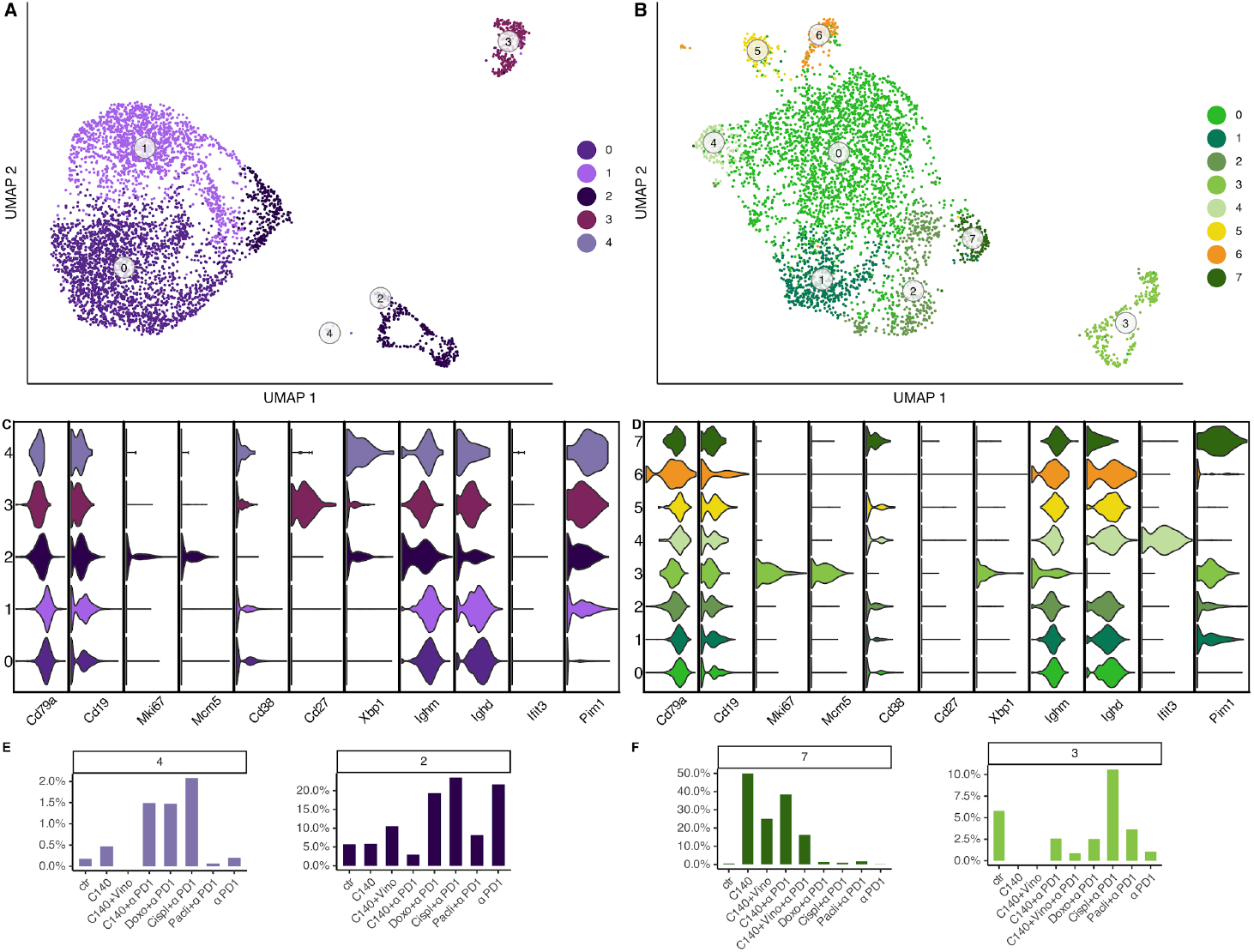
B-like cell sub-clustering analysis in 4T1 and EMT6.

UMAP of B cell sub-clustering in 4T1 (A) and EMT6 (B). Gene signature of B cell sub-clustering in 4T1 (C) and EMT6 (D). Proportions of cells in relevant 4T1 (E) and EMT6 (F) B cell sub-clusters among different conditions.

Strikingly, the proportions of cells sharing the same transcriptional profile were generally comparable between the treated and untreated TMEs of the two cell lines. Indeed the germinal B cells (cluster 4 and 7, in 4T1 and EMT6 respectively) in the untreated sample were few while in the C140+αPD-1 treatments increased in both the murine models (Fig. 5F-H). In addition, similarities on the proportions of the proliferative B cell clusters were found. Specifically, cluster 2, for 4T1, decreased only in correspondence of cyclophosphamide alone or in combination with the immunotherapy compared to the control; while the percentage of cells in cluster 3 in EMT6 decreased in all the treatments but the cisplatin in combination with anti-PD-1 (Fig. 5E, G). Interestingly, Cispl+αPD-1 condition presented the highest percentage of proliferative B cells for both the clusters, suggesting a common behaviour of the two tumours in response to this specific combination of treatment (Fig. 5C, fig. S7).

## Discussion

Although recently, many studies focused on the immune TME of BC ^47, 60, 63, 64^, specific knowledge about their immune cell composition is still limited. Here we report a fine characterization of the immune transcriptional profiles of almost 50,000 single cells in two TNBC murine cell line TMEs. Major differences across the two cell line TMEs were detected; these included a major macrophagic component in the EMT6 TME while a great number of Cd4- and Cd8-like T cells into 4T1 TME. Finally, a comparable percentage of B cells and neutrophils were observed in both the tumoral models. This is in agreement with results based on flow cytometry that detected a large infiltration of Cd4^+^ and Cd8^+^ in the TME of 4T1 cell line, whereas the myeloid cells represent a minor component ^26^. However, few points should be carefully considered when comparing scRNA-seq and flow cytometry technologies: the first is related to the lower number of cells that the scRNA-seq analyses have in comparison to classic flow cytometry; the second is related to the challenge of comparing populations previously characterised with surface protein markers (using flow cytometry), and for which a gene expression profile is unknown, with cell populations finely characterised by only gene expression profiles (using scRNA-seq technique); the third is related to different laboratory procedures and/or operators that might influence the number of isolated cells.

Sub-clustering of T cells in the TMEs of the 4T1 cell line revealed previously uncharacterised sub-populations with unique transcriptional profiles. Trajectory analyses highlighted some of these populations as intermediate states and identified a tripartite differentiation for both Cd4- and Cd8-like T cells as also reported in human TNBC TMEs ^53^. Interestingly, percentages of cells in these sub-clusters varied significantly across the conditions and in particular, subsets associated with regulatory and γδ T cells (expressing high level of *Il17a*), were found decreasing in conditions with a higher pre-clinical efficacy in *in vivo* experiments ^27^. Although the association of regulatory T cells and poor prognosis in multiple cancer types has been widely characterized ^65–69^, the pro-tumoral activity of γδ T cells in murine TNBC was poorly or never observed at scRNA-seq level ^70^, therefore, here we helped in validating their function along with the expression of *Il17a* ^49,55^ and pave the way to a further gene signature classification of this specific subset. Furthermore, we confirmed the previously reported general increase of exhausted-like Cd8 T cell subpopulation in pre-clinical treatments with low *in vivo* efficacy and in the untreated samples ^27^. This is in line with the molecular mechanism by which tumours escape the immune system exploiting checkpoint inhibitors, whose persistent activation induces a state of exhaustion among T cells ^19^. On the other hand, a proliferative Cd8-like T cell subcluster (expressing high levels of *Mki67*) was found to increase in correspondence of the treatments with cyclophosphamide alone or in combination with other chemotherapy/immunotherapy. This is in accordance with the association of proliferative Cd8 T cells and better outcome in cancer ^71^.

Macrophage-like cells on the TMEs of EMT6 TNBC revealed subtypes expressing genes related to M2-like macrophages enriched in untreated condition and in treatments with poor efficacy in *in vivo* experiments ^27^. This confirms their known activity in favour of tumour progression and metastasization ^72^. Contrarily, clusters expressing genes related to M1-like macrophages were found enriched in TMEs corresponding to high efficacy preclinical treatments.

Finally, we suggested a common behaviour, along some conditions (C140+αPD-1, C140+V and Cispl+αPD-1), of two B cell sub-clusters presenting similar gene signatures in both the murine tumor cell lines. These clusters were associated with proliferative B and germinal B cells and followed opposite trends for some of the high efficacy treatments. In particular the alkylating agent cisplatin in combination with the immunotherapy αPD-1 favoured the expansion of germinal B cells. Moreover, plasma B cells, only identified in 4T1 TME, increased in high efficacy treatments such as cyclophosphamide in combination with other chemotherapies or anti-PD-1 (Fig. S7), confirming their association with improved survival ^73,74^. However, a cross investigation, also considering the tertiary lymphoid structures, is needed to clarify the characterization and response of these B cell populations.

In summary, the identification of unknown and cancer-resistant immune cell populations after tumour removal and the fine-scale characterization of the immune TME in this work could be an important resource for the diagnosis and treatment of TNBC leading to an application at clinical level.

## Future perspectives and limitations of the study

In this context a detailed extension and validation of the analyses is required due to some limitations related to scRNA-seq. In particular, the number of cells in some sub-clusters were really low and therefore a more detailed analyses either using classic flow cytometry or a scRNA seq only on sorted cells belonging to those specific sub-clusters is required. In addition, more replications of each condition reported in this work and the transcriptional investigation of CD45^−^ cells populating the two TMEs with the aim to investigate also the release of specific chemokines and cytokines of the tumour cell could strengthen the results obtained here.

Moreover, although scRNA seq is certainly a powerful technology, it is a single profiling strategy with a limited detection efficiency for key markers, as mentioned above ^75^. This issue can be addressed by recent technologies that incorporate oligonucleotide-labelled antibodies into droplet-based scRNA seq to measure at the same time, surface markers alongside intracellular mRNA transcripts. For example cellular indexing of transcriptomes and epitopes by sequencing (CITE-seq) provides an additional layer of information for the same cell by combining both transcriptomics with proteomics data and reduces the overall cost of high-throughput sequencing on multiple samples ^76^.

## Data Availability

Raw count matrices generated with the scRNA-seq Sankemake CellRanger v4 alignment pipeline and pre-QC Seurat objects are available under the under accession number GSEXXXXX at the GEO (http://www.ncbi.nlm.nih.gov/geo/).

## Codes Availability

The Snakemake CellRanger v4 pipeline is available at this link https://github.com/raveancic/scRNAaltas_TNBC_mm/tree/master/cl_crt_FASTQ2countmat

## Acknowledgements

This research was funded by AIRC (IG20109) and the Italian Ministry of Health (Ricerca Corrente). L.C. is a PhD student within the European School of Molecular Medicine (SEMM). The authors would like to thank Stefano Cheloni for fruitful suggestions during the first step of the alignment analyses; Michel Gerard Arnaud Ceol for computational support.

## Author information

F.B directed the study. F.B and A.R designed the study. L.C performed the analyses under the supervision of A.R and F.B. S.M and R.H provided computational support for the analyses. P.F., S.O., G.M and P.M provided support for the interpretation of laboratory analyses. A.R, L.C and F.B wrote the original draft. All the authors discussed the results and contributed to the final version of the manuscript.

## Supplementary Figure Legend

**Fig. S1. Single-cell RNA seq quality controls of two tumoral models**. Number of UMI plotted on x-axis versus number of genes on y-axis divided by condition for 4T1 (A) and EMT6 (B). Each dot represents a cell colour-coded by the mito ratio. Darker colours mean low mitochondrial expression, lighter colours indicate high mitochondrial genes. Vertical and horizontal red bars indicate the threshold used for number of genes (y-axis) and number of transcripts (x-axis).

**Fig. S2. Batch effect on 4T1 cell line**. A) UMAP showing batch effect on 4T1 cell line. Each cell is represented by a colored dot colored according to the 10X chemistry used. B) UMAP showing batch effect correction on 4T1. (Orange for V2 and blue for V3).

**Fig. S3. Cell Assignation**. Automatic cell populations assignment for 4T1 cell line (A) and EMT6 cell line (B). Clusters are arranged to the x-axis while the percentage of cells are signed on the y-axis. The colors are referring to the different types of cells automatically assigned by SingleR. C) UMAPs of known marker gene expression in 4T1 cell line. *Cd3e*, *Cd4*, *Cd8b1* genes were evaluated for T cells; *Ncr1* for NK cells; *Cd19* for B cells; *Csf3r* for neutrophils, *Cd68* for macrophages; *Basp1* for DC. Each dot represents a cell. The intensity of the colour (from yellow to dark violet) indicates the greatest expression of the specific marker gene. D) UMAP of known marker gene expression in EMT6 cell line. *Cd3e*, *Cd4*, *Cd8b1* genes were evaluated for T cells; *Ncr1* for NK cells; *Cd19* for B cells;*Csf3r* for neutrophils, *Adgre1* for macrophages; *Basp1* for DC. Each dot represents a cell. The intensity of the colour (from yellow to dark violet) indicates the greatest expression of the specific marker gene.

**Fig. S4. Cd4-like T cell sub-cluster variation**.Percentage of cells in each 4T1 Cd4-like T cell sub-cluster among different conditions. Bar graph shows on the x-axis the conditions, while the percentage of cells per cluster is plotted in the y-axis.

**Fig. S5. Cd8-like T cell sub-cluster variation**.Percentage of cells in each 4T1 Cd8-like T cell sub-cluster among different conditions. Bar graph shows on the x-axis the conditions, while the percentage of cells per cluster is plotted in the y-axis.

**Fig. S6. Macrophage sub-cluster variation**.Percentage of cells in each EMT6 macrophages sub-cluster among different conditions. Bar graph shows on the x-axis the conditions, while the percentage of cells per cluster is plotted in the y-axis.

**Fig. S7. B cell sub-cluster variation**.Percentage of cells in B cell sub-clusters for 4T1 (A) and EMT6 (B) cell lines among different conditions. Bar graph shows on the x-axis the conditions, while the percentage of cells per cluster is plotted in the y-axis.

## Table legend

**Table 1. Condition overview**.Schematic view of conditions (samples) with the corresponding treatment for the 4T1 and EMT6 cell line TMEs with the 10x Genomic chemistry version used for each condition.

## Supplementary Table Legend

**Table S1**. Top 20 differentially expressed genes for the 4T1 cell line

**Table S2**. Top 20 differentially expressed genes for the EMT6 cell line

**Table S3**. Number of cells pre- and post-quality control for each condition in 4T1 cell line

**Table S4**. Number of cells pre- and post-quality control for each condition in EMT6 cell line

