## Supplementary Figures for "A single-cell transcriptomic landscape of innate and adaptive intratumoral immunity in triple negative breast cancer during chemo- and immunotherapies"

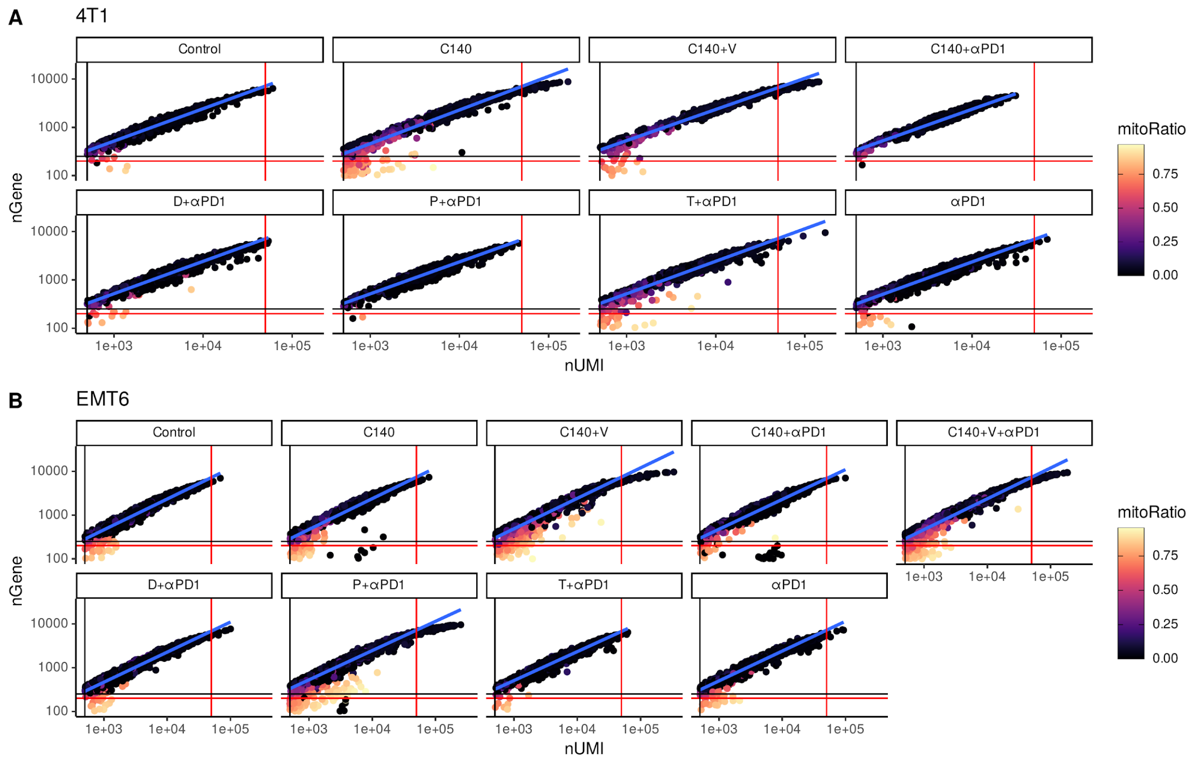


**Fig. S1. Single-cell RNA seq quality controls of two tumoral models.** Number of UMI plotted on x-axis versus number of genes on y-axis divided by condition for 4T1 (A) and EMT6 (B). Each dot represents a cell colour-coded by the mito ratio. Darker colours mean low mitochondrial expression, lighter colours indicate high mitochondrial genes. Vertical and horizontal red bars indicate the threshold used for number of genes (y-axis) and number of transcripts (x-axis).

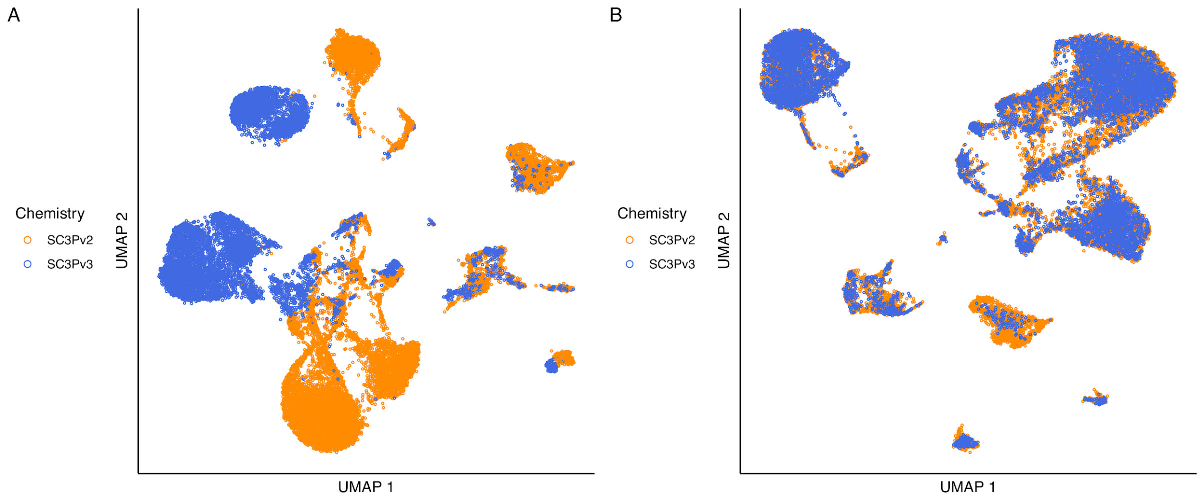


**Fig. S2. Batch effect on 4T1 cell line.** A) UMAP showing batch effect on 4T1 cell line. Each cell is represented by a colored dot colored according to the 10X chemistry used. B) UMAP showing batch effect correction on 4T1. (Orange for V2 and blue for V3).

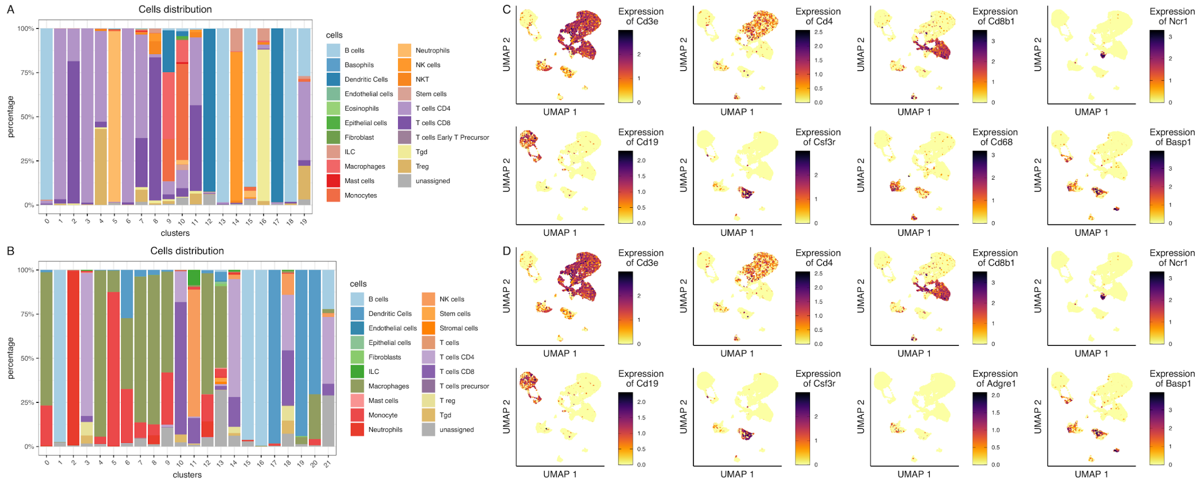


**Fig. S3. Cell Assignation.** Automatic cell populations assignment for 4T1 cell line (A) and EMT6 cell line (B). Clusters are arranged to the x-axis while the percentage of cells are signed on the y-axis. The colors are referring to the different types of cells automatically assigned by SingleR. C) UMAPs of known marker gene expression in 4T1 cell line. Cd3e, Cd4, Cd8b1 genes were evaluated for T cells; Ncr1 for NK cells; Cd19 for B cells; Csf3r for neutrophils, Cd68 for macrophages; Basp1 for DC. Each dot represents a cell. The intensity of the colour (from yellow to dark violet) indicates the greatest expression of the specific marker gene. D) UMAP of known marker gene expression in EMT6 cell line. Cd3e, Cd4, Cd8b1 genes were evaluated for T cells; Ncr1 for NK cells; Cd19 for B cells; Csf3r for neutrophils, Adgre1 for macrophages; Basp1 for DC. Each dot represents a cell. The intensity of the colour (from yellow to dark violet) indicates the greatest expression of the specific marker gene.

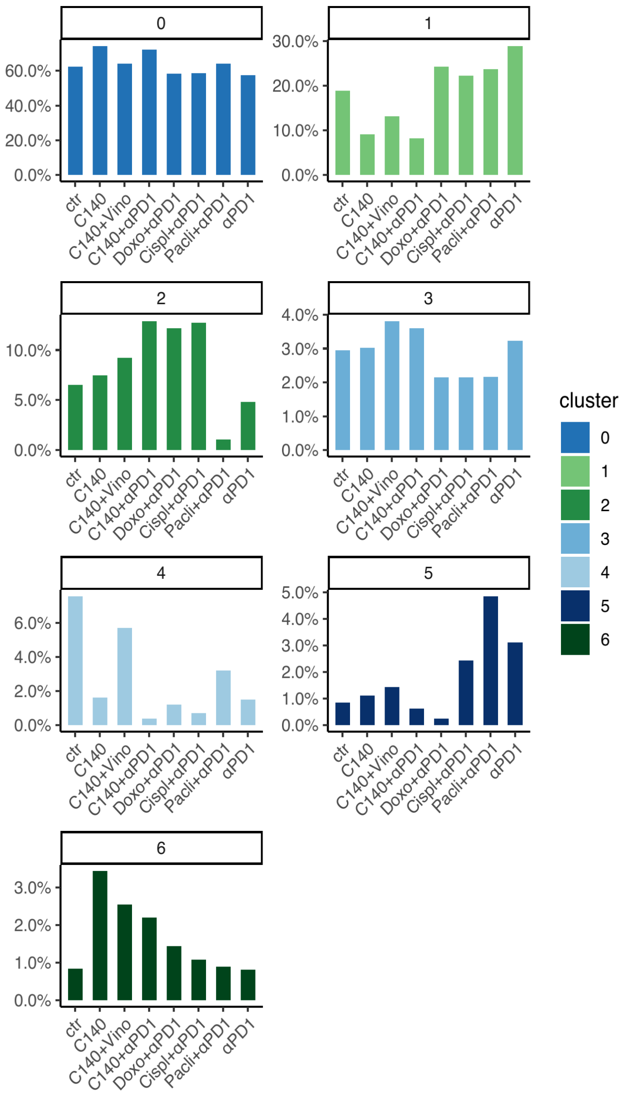


**Fig. S4. Cd4-like T cell sub-cluster variation.** Percentage of cells in each 4T1 Cd4-like T cell sub-cluster among different conditions. Bar graph shows on the x-axis the conditions, while the percentage of cells per cluster is plotted in the y-axis.

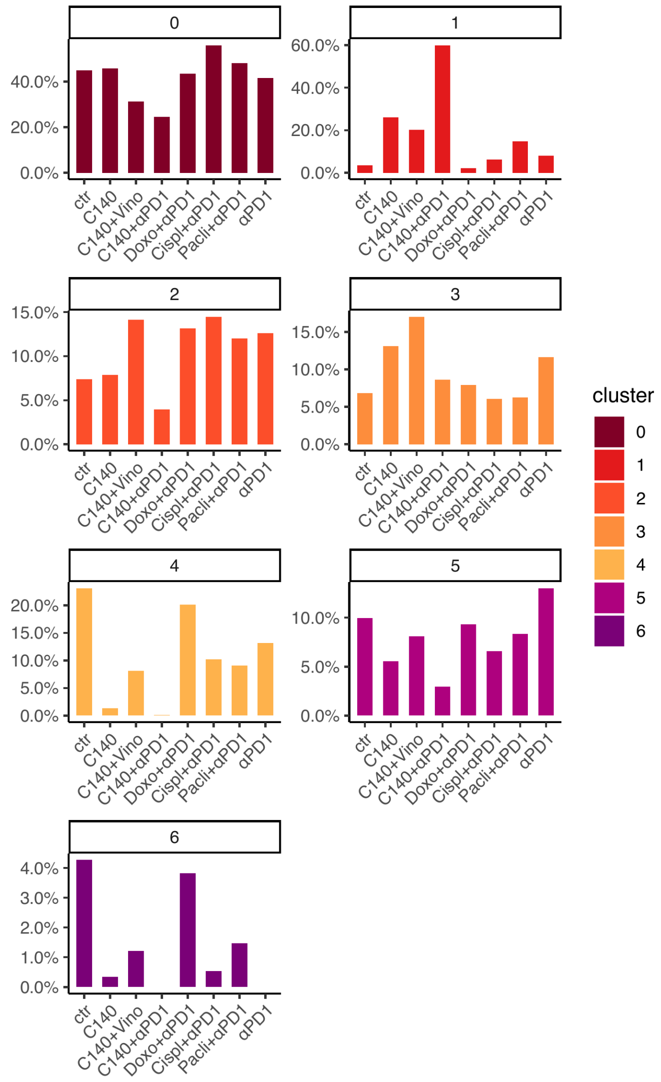


**Fig. S5. Cd8-like T cell sub-cluster variation.** Percentage of cells in each 4T1 Cd8-like T cell sub-cluster among different conditions. Bar graph shows on the x-axis the conditions, while the percentage of cells per cluster is plotted in the y-axis.

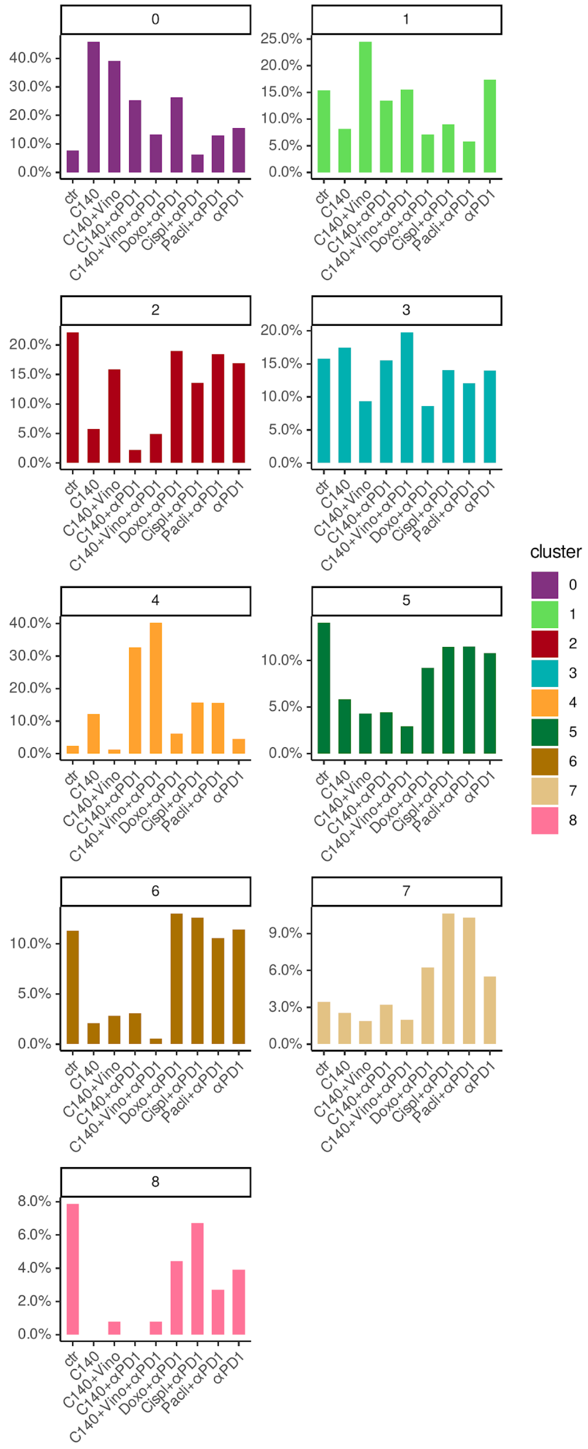


**Fig. S6. Macrophage sub-cluster variation.** Percentage of cells in each EMT6 macrophages sub-cluster among different conditions. Bar graph shows on the x-axis the conditions, while the percentage of cells per cluster is plotted in the y-axis.

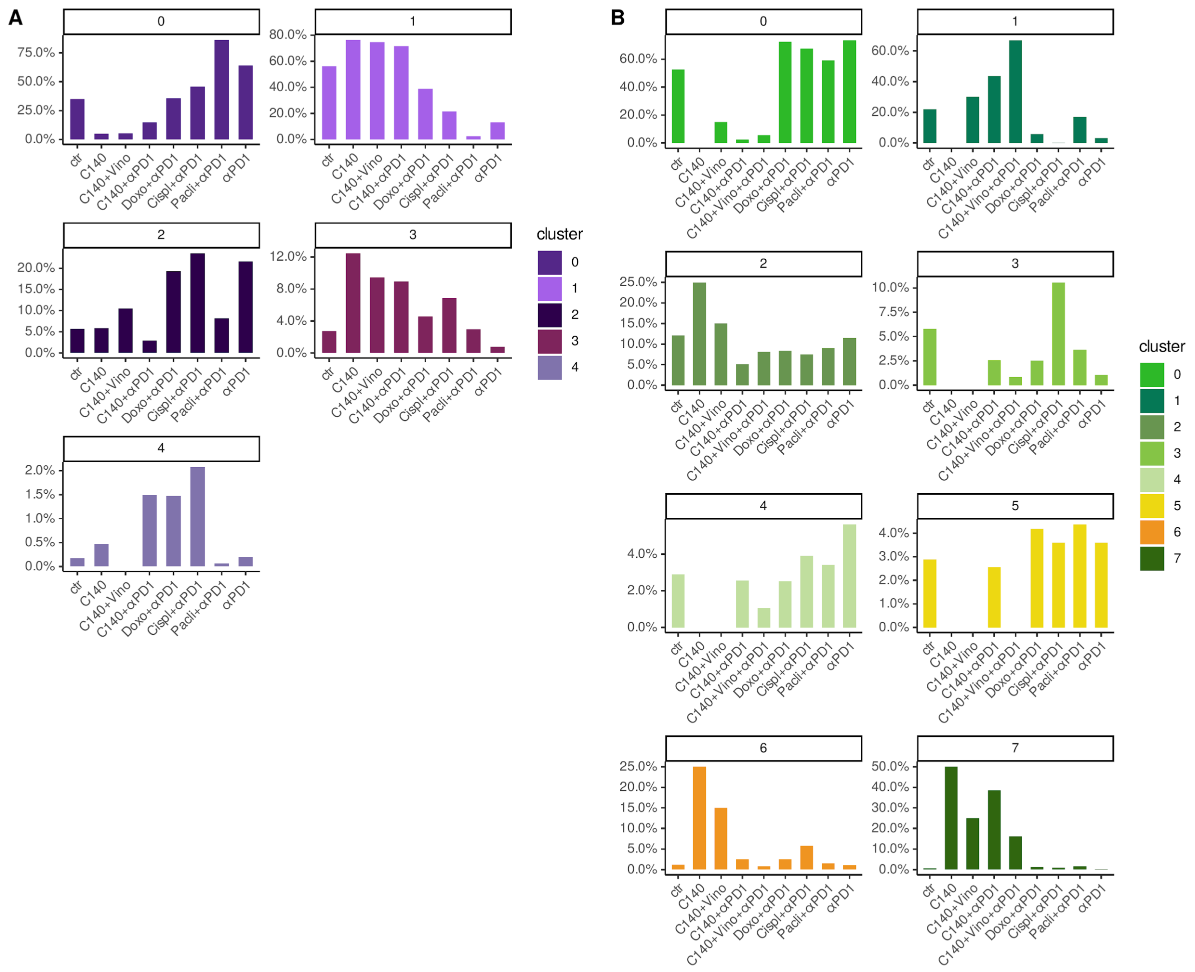


**Fig. S7. B cell sub-cluster variation**. Percentage of cells in B cell sub-clusters for 4T1 (A) and EMT6 (B) cell lines among different conditions. Bar graph shows on the x-axis the conditions, while the percentage of cells per cluster is plotted in the y-axis.
